# Consequences of Early Postnatal Blockade of Aldosterone Synthesis on Behaviour and Stress Response in Male and Female Rats

**DOI:** 10.64898/2026.07.28.741170

**Authors:** L. Karailievova, P. Karailiev, A. Nagyova, D. Jezova, N. Hlavacova

## Abstract

**Aim:** The aim of the present study was to determine whether pharmacological inhibition of aldosterone synthesis during the stress-hyporesponsive period (SHRP) affects behaviour and adrenocortical stress responsiveness later in development and whether these effects differ between males and females.

**Methods:** Newborn Wistar rat pups (males n=40, females n=40) were treated with aldosterone synthase inhibitor FAD286 (30 mg/kg per day, orally) or vehicle from PND3 to PND9. To verify the pharmacodynamic action of FAD286, serum and adrenal glands from 10-day-old pups were analysed. The remaining pups were weaned on PND21 and underwent open-field (PND23), elevated plus-maze (PND29) and salt-preference testing. At PND46, half of each group was exposed to restraint stress for 120 min.

**Results:** In 10-day-old pups, treatment with FAD286 resulted in increased gene expression of CYP11B2 (aldosterone synthase) and CYP11B1 (11-beta-hydroxylase) in the adrenal glands, increased serum levels of corticosterone, and decreased concentrations of serum aldosterone. FAD286 did not modify the general locomotor activity assessed in juvenile rats. Inhibition of aldosterone synthase by FAD286 resulted in altered anxiety-like behaviour in a sex-dependent manner. Postnatal FAD286 treatment led to increased anxiety-like behaviour in female, but not male rats. During adolescence, early FAD286 treatment increased overall aldosterone concentrations without altering the aldosterone response to restraint. Basal corticosterone concentrations were unchanged, whereas the response to restraint was enhanced.

**Conclusions:** The present study demonstrates that transient inhibition of aldosterone synthesis during the SHRP led to alterations in anxiety-related behaviour and adrenocortical regulation later in development, with some behavioural effects being sex-dependent.

## 1. Introduction

The hypothalamic–pituitary–adrenocortical (HPA) axis is a central component of the neuroendocrine response to stress. Activation of the HPA axis stimulates glucocorticoid release from the adrenal cortex, thereby maintaining homeostasis and facilitating adaptation to stressors (Herman et al., 2016). Although glucocorticoids are essential for an adequate stress response, excessive or prolonged glucocorticoid exposure may adversely affect neuronal maturation and brain plasticity, particularly during early development (Welberg and Seckl, 2001; Lupien et al., 2009). In rats, early postnatal development is characterized by the stress-hyporesponsive period (SHRP), during which basal corticosterone concentrations are low, and corticosterone responses to several stress stimuli are markedly attenuated (Sapolsky and Meaney, 1986; Levine, 2001; Schmidt, 2019). This reduced glucocorticoid responsiveness is thought to protect the developing brain from excessive exposure to glucocorticoids. Importantly, the SHRP does not represent a general suppression of adrenocortical responsiveness.

The response of other adrenocortical hormones during this developmental period has received much less attention. Aldosterone, the principal mineralocorticoid hormone of the adrenal cortex, is secreted not only in response to changes in fluid and electrolyte balance but also during stress exposure (Sowers et al., 1981; Stier et al., 2004). Its secretion is regulated mainly by the renin–angiotensin system and plasma potassium concentrations, although adrenocorticotropic hormone may contribute to the aldosterone response during acute stress. We have previously demonstrated that 10-day-old rat pups exhibit a pronounced increase in plasma aldosterone concentrations in response to several acute stress stimuli, despite their markedly reduced corticosterone response (Varga et al., 2013). Moreover, the stress-induced aldosterone response was greater in pups than in adult rats. These findings suggest that aldosterone may represent an important adrenocortical stress hormone during the SHRP, when corticosterone responsiveness is markedly reduced.

Aldosterone has traditionally been studied primarily for its role in the regulation of water and electrolyte balance and cardiovascular function (Hattangady et al., 2012). Nevertheless, increasing evidence indicates that its actions are also relevant to stress-related behaviour and affective disorders. Our previous studies in adult rats have demonstrated that prolonged elevation of circulating aldosterone induces anxiety- and depression-like behaviour (Hlavacova and Jezova, 2008; Hlavacova et al., 2012) and aldosterone signals the onset of depressive behaviour in an animal model of treatment-resistant depression (Franklin et al., 2015). Systemic treatment with the mineralocorticoid receptor antagonist eplerenone produced an anxiolytic profile accompanied by changes in stress hormone release (Hlavacova et al., 2010). Anxiolytic-like effects have also been observed following central corticosteroid receptor blockade (Korte et al., 1995; Bitran et al., 1998). Clinical findings also suggest an association between aldosterone and major depressive disorder. Salivary aldosterone concentrations have been associated with treatment outcome (Büttner et al., 2015) and with the duration and severity of depressive episodes (Segeda et al., 2017; Izakova et al., 2020). These observations support the potential relevance of aldosterone signalling to the pathophysiology of affective disorders.

Biological sex is an important determinant of stress responsiveness. Sex differences have been identified at several levels of the HPA axis, including the central regulation of corticotropin-releasing hormone secretion, pituitary responsiveness, adrenal glucocorticoid production and feedback regulation. These differences are influenced by gonadal hormones, developmental and organisational mechanisms and may change across the lifespan. Importantly, the magnitude and even the direction of sex differences depend on the developmental stage, the nature and duration of the stressor and the parameters being evaluated (Goel et al., 2014; Oyola and Handa, 2017; Bangasser and Wiersielis, 2018; Heck and Handa, 2019). Similarly, experimental manipulations during early life may produce different behavioural and neuroendocrine consequences in males and females. Sex should therefore be considered when evaluating the long-term behavioural and neuroendocrine consequences of endocrine manipulations performed during sensitive developmental periods.

Experimental evidence on the behavioural and neuroendocrine effects of aldosterone has been obtained predominantly in adult animals. Its physiological role during early postnatal development has received little attention, although the regulation of adrenocortical stress responses during this period differs substantially from that in adulthood. In particular, it remains unknown whether the pronounced aldosterone response observed during the SHRP contributes to the subsequent development of behavioural and neuroendocrine stress regulation. The long-term consequences of transient pharmacological inhibition of aldosterone synthesis during this developmental period have not yet been investigated.

The aim of the present study was to determine whether pharmacological inhibition of aldosterone synthesis during the stress-hyporesponsive period affects behaviour and adrenocortical stress responsiveness later in juvenile and adolescent periods. Aldosterone synthase was inhibited by the selective inhibitor, FAD286, during early postnatal development, and the later behavioural and neuroendocrine consequences were evaluated in both sexes. We hypothesised that reduced aldosterone availability during this developmental period would result in persistent alterations in behavioural and neuroendocrine responses to stress and that these effects might differ between males and females.

## 2. Materials and Methods

### 2.1. Animals

A total of nine adult timed-pregnant Wistar rats (AnLab, Prague, Czech Republic) arrived at the animal facility on gestational day 16. Approximately one week later, the litters were born, and the day of birth was designated as postnatal day 0 (PND0). On PND3, litters were adjusted to eight or nine pups per dam to reduce competition for maternal milk during the early postnatal period. A total of 80 pups, 40 males and 40 females, were included in the study. Pups retained for the long-term part of the study remained with their dams until weaning on PND21. Thereafter, animals were housed two or three per cage according to sex and treatment. Rats were kept under standard housing conditions with a constant 12:12 h light/dark cycle (lights on at 06:00 h), a temperature of 22 ± 2 °C, and a relative humidity of 55 ± 10%. Food and water were available ad libitum. All experimental procedures were approved by the Animal Health and Animal Welfare Division of the State Veterinary and Food Administration of the Slovak Republic (Approval No. 2876-3/2020-220) and conformed to the EU Directive 2010/63/EU on the protection of animals used for scientific purposes.

### 2.2. Study design and FAD286 treatment

The overall experimental design is shown in Fig. 1. On PND3, pups were sexed and assigned to treatment with the aldosterone synthase inhibitor FAD286 or vehicle (VEH). The study included 40 males and 40 females, with 20 animals/sex assigned to each treatment condition. Within each litter, pups were distributed as evenly as possible between the vehicle and FAD286 treatment groups and, depending on the sex composition of the litter, across the four sex × treatment groups.

**Figure 1.**
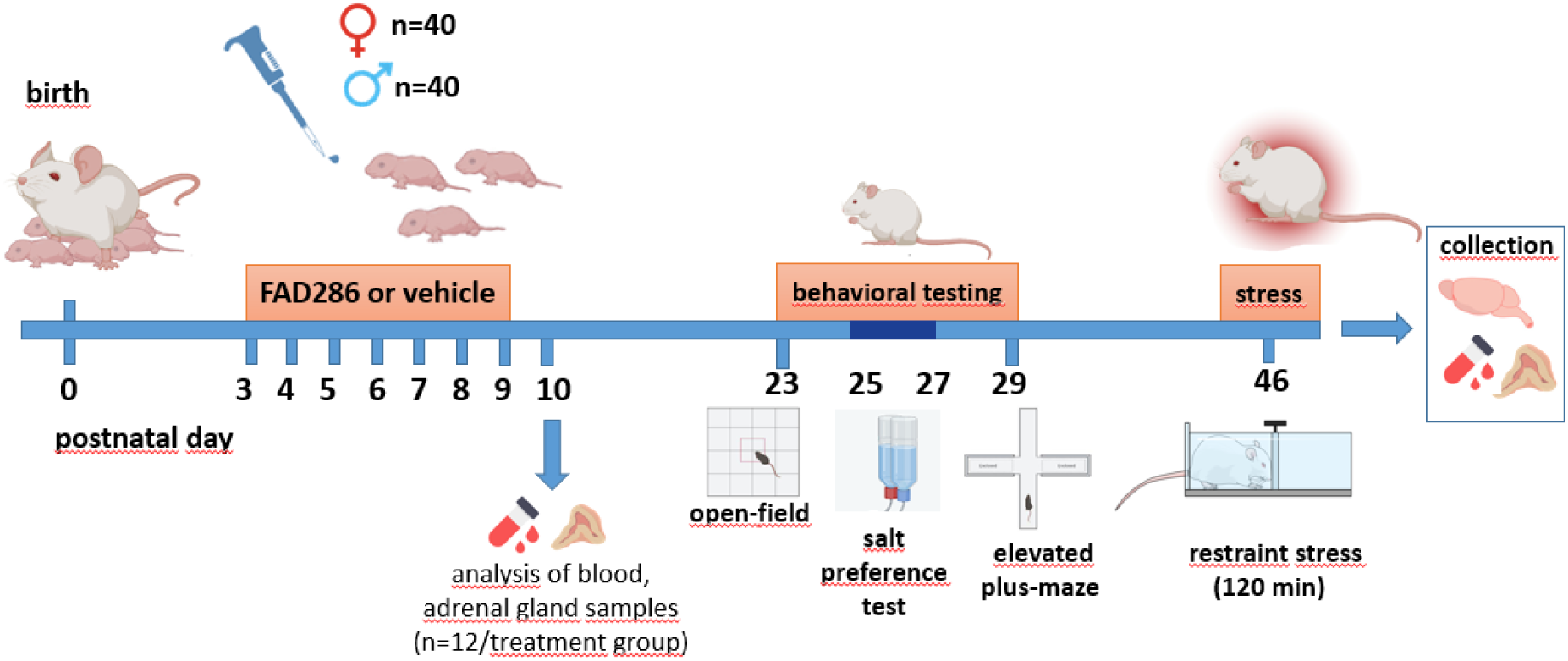
Schematic overview of the experimental design. Male and female rat pups were treated orally with the aldosterone synthase inhibitor FAD286 or vehicle from postnatal day (PND) 3 to PND9. On PND10, a subset of animals from each treatment group was killed for blood and adrenal gland collection (n = 12 per treatment group). The remaining animals underwent behavioural testing in the open-field test on PND23 and the elevated plus-maze test on PND29. Salt preference test was performed on PND25-27. On PND46, approximately half of the animals from each sex and treatment group were exposed to 120 min of restraint stress, followed by sample collection.

FAD286 (Sigma-Aldrich, USA) was dissolved in distilled water and administered orally at a dose of 30 mg/kg body weight once daily from PND3 to PND9. The dose of FAD286 was selected based on previous studies (Menard et al., 2006; Deliyanti et al., 2012). Vehicle-treated pups received an equivalent volume of distilled water. FAD286 and vehicle were delivered directly into the oral cavity using a micropipette at approximately 14:00 h. The individual dose was adjusted according to body weight.

During the treatment period, the surface righting reflex was assessed on PND6. On PND10, 12 animals from each treatment group were killed by decapitation, with males and females approximately equally represented in both groups. Blood and adrenal glands were collected for measurements of circulating aldosterone and corticosterone concentrations and adrenal expression of steroidogenic enzymes.

The remaining 56 pups were kept with their dams until weaning on PND21 and were assigned to four experimental groups: VEH-treated females, FAD286-treated females, VEH-treated males, and FAD286-treated males (n = 14 per group). After weaning, animals were housed two or three per cage according to sex and treatment and underwent behavioural testing during the juvenile period. General locomotor activity and anxiety-like behaviour were assessed in the open-field test on PND23, salt preference was evaluated from PND25 to PND27, and anxiety-like and risk-assessment behaviour was assessed in the elevated plus-maze test on PND29. On PND46, half of the animals from each experimental group (n=7/group) were exposed to restraint stress for 120 min, whereas the remaining animals served as non-stressed controls. Immediately after the stress procedure, stressed animals and the corresponding non-stressed controls were killed by decapitation.

### 2.3. Behavioural testing

#### 2.3.1. Surface righting reflex

The surface righting reflex was assessed on PND6, as described previously (Naik et al., 2015). Each pup was placed in a supine position on a flat surface, and the presence or absence of the righting response was recorded. A positive response was defined as returning to a prone position with all four paws in contact with the surface.

#### 2.3.2. Open-field test

On PND23, general locomotor activity and anxiety-like behaviour were assessed in the open-field test, as described previously with minor modifications (Hlavacova et al., 2023). The apparatus consisted of a square arena measuring 60 × 60 cm, surrounded by 60-cm-high walls. The central area was defined as a square measuring 20 × 20 cm. At the beginning of the test, each animal was gently placed in the corner of the open-field arena, facing the wall, and allowed to explore the arena freely for 10 min. The apparatus was illuminated with dim light at 30 lx in the central area and 20 lx in the peripheral area. Testing was performed during the light phase of the light/dark cycle between 13.00 h and 15.00 h. General locomotor activity was evaluated by recording the total distance travelled. Anxiety-like behaviour was assessed by the number of entries into and the time spent in the central area of the arena. Behavioural parameters were analysed using EthoVision XT 9.0 software (Noldus Information Technology, Wageningen, the Netherlands).

#### 2.3.3. Salt Preference Test

Salt preference was assessed using a two-bottle choice test, as described previously (Flynn et al., 1993; Inui-Yamamoto et al., 2017). The test was conducted from PND25 to PND27. Rats remained group-housed, with two or three animals of the same sex and treatment group per cage. During the 48-h testing period, each cage was provided with two bottles, one containing 0.3 M NaCl solution and the other containing tap water. Food and fluids were available ad libitum, and no water or sodium deprivation was applied before or during the test. To minimise potential effects of side preference, the positions of the bottles were exchanged after 24 h.

The bottles were weighed at the beginning of the test and after each 24-h period. Consumption of the 0.3 M NaCl solution and tap water was calculated from the corresponding changes in bottle weight and expressed per cage. Mean daily fluid intake was calculated by averaging the values recorded during the two consecutive 24-h periods. Relative NaCl solution intake was calculated as grams per 100 g of the combined body weight of the animals housed in the respective cage and expressed as grams per 100 g of body weight. Salt preference was calculated as the consumption of a 0.3 M NaCl solution divided by total fluid consumption and expressed as a percentage: Salt preference (%) = NaCl solution intake / (NaCl solution intake + water intake) × 100. As fluid consumption was measured collectively, the cage was considered the experimental unit.

#### 2.3.4. Elevated plus-maze test

On PND29, anxiety-like behaviour was assessed in the elevated plus-maze test, as described previously (Hlavacova et al., 2010; Chmelova et al., 2019). The apparatus consisted of two opposite open arms (50 × 10 cm) and two opposite closed arms (50 × 10 × 40 cm) extending from a central platform (10 × 10 cm) to form a plus-shaped maze. The maze was elevated 50 cm above the floor. At the beginning of the test, each animal was placed on the central platform facing an open arm and was allowed to explore the maze freely for 5 min. The apparatus was illuminated with dim light at 45 lx in the open arms and 15 lx in the closed arms. Testing was performed during the light phase of the light/dark cycle between 13.00 h and 15.00 h. Spatiotemporal measures included the number of entries into and the percentage of time spent in the open and closed arms. The number of open-arm entries and the percentage of time spent in the open arms were used as measures of anxiety-like behaviour. An arm entry was defined as all four paws entering the respective arm. Ethological parameters related to exploration and risk-assessment behaviour were also evaluated, including head dipping and stretched-attend postures. Head dipping was defined as an exploratory movement in which the animal protruded its head over the side of an open arm and down towards the floor. A stretched-attend posture was defined as an exploratory posture in which the animal stretched forward and subsequently returned to its original position without moving forward. The total frequencies of head dips and stretched-attend postures were used for analysis. Spatiotemporal parameters were analysed using EthoVision XT 9.0 software (Noldus Information Technology, Wageningen, the Netherlands). Ethological parameters were scored manually from video recordings by a trained observer blinded to the experimental conditions.

### 2.4. Restraint stress

On PND46, seven animals from each sex and treatment group were exposed to restraint stress for 120 min. Rats were placed individually in transparent cylindrical restrainers adjusted to their body size. The restrainers prevented turning around and markedly restricted locomotion without compressing the body. The restraint apparatus was of the type described previously (Chomanic et al., 2022). Non-stressed control animals remained undisturbed in their home cages for the corresponding period. Immediately after the restraint period, stressed animals and the corresponding non-stressed controls were killed by decapitation, and blood and adrenal gland samples were collected.

### 2.5. Tissue and blood collection

Immediately after decapitation, trunk blood was collected into polyethene tubes without an anticoagulant. The blood was centrifuged at 3000 rpm for 15 min at 4 °C, and the serum was separated and stored at -30 °C until analysis. The right and left adrenal glands, as well as the thymus, were quickly removed, weighed, frozen in liquid nitrogen, and stored at -70 °C until analysed.

### 2.6. Hormone measurements

Serum aldosterone concentrations were measured by a radioimmunoassay (RIA) using a commercially available kit (Aldosterone, RIA CT, 96 tests, R-CW-100, DIAsource, Belgium). The intra- and inter-assay coefficients of variation were 3.8% and 7.5%, respectively. The detection limit was 1.4 pg/ml. Serum corticosterone concentrations were measured using a commercially available RIA kit (rCorticosterone [125 I] RIA KIT, RK-548CT, Institute of Isotopes Co. Ltd., Hungary) according to the manufacturer’s instructions. The assay’s detection limit was 0.77 μg/100 mL. The corticosterone assay had intra- and inter-assay coefficients of variation of 4.4% and 7.2%, respectively. All samples were analysed in duplicate.

### 2.7. RNA isolation and quantitative real-time PCR analysis

Adrenal mRNA expression of *Cyp11b2*, encoding aldosterone synthase, and *Cyp11b1,* encoding steroid 11β-hydroxylase, was measured in left adrenal gland samples collected on PND10 and PND46 using quantitative real-time PCR. Total RNA was extracted using TRI Reagent® (Sigma-Aldrich, Merck KGaA, Germany) according to the manufacturer’s instructions. One μg of total RNA was reverse-transcribed into cDNA using oligo(dT) primers and the ProtoScript® First Strand cDNA Synthesis Kit (New England Biolabs, USA). Quantitative real-time PCR was performed in a final reaction volume of 10 μL using Luna® Universal qPCR Master Mix (New England Biolabs, Ipswich, MA, USA) on a QuantStudio™ 5 Fast Real-Time PCR System (Applied Biosystems, Thermo Fisher Scientific, USA). Gene-specific primers were used at a final concentration of 0.25 pmol/μL, and 5 ng of cDNA was added to each reaction. Primer sequences are listed in Table 1. Primers were designed using NCBI Primer-BLAST. Amplification specificity was verified by melting-curve analysis. The quantity of each transcript was determined from a gene-specific standard curve. Expression of the target genes was normalised to the geometric mean of the quantities of peptidylprolyl isomerase A *(Ppia*) and ribosomal protein S29 (*Rps29*). Normalised gene expression was calculated as the ratio of the target-gene quantity to the geometric mean of the two reference-gene quantities and expressed in arbitrary units.

**Table 1.**
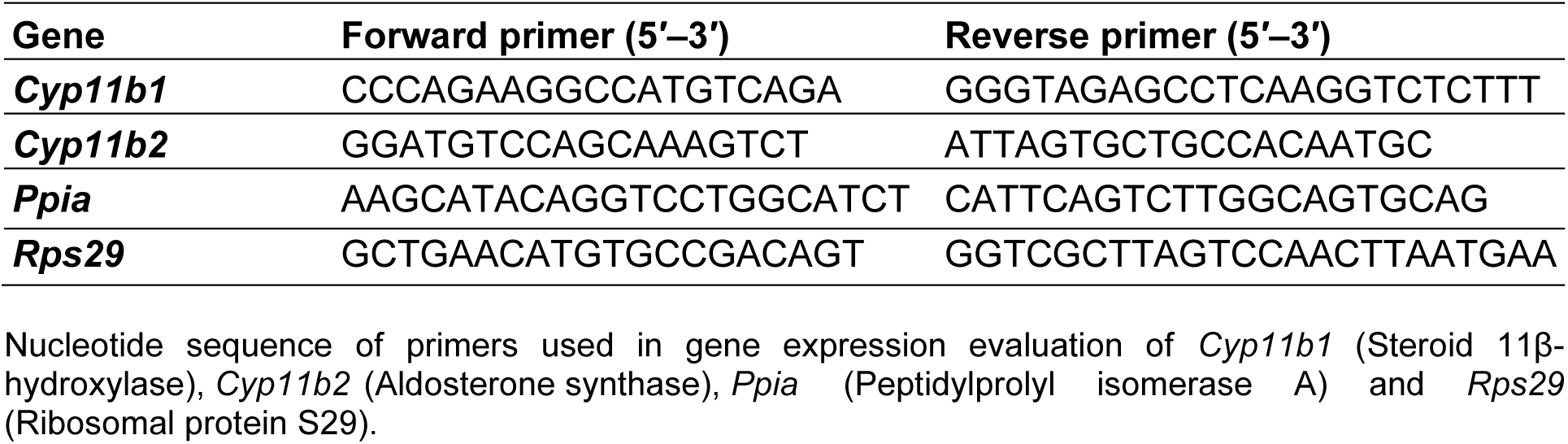
Primer sequences used for quantitative real-time PCR.

### 2.8. Statistical analysis

Data were assessed for normality using the Shapiro–Wilk test. The surface righting reflex was recorded as a binary outcome and summarised descriptively. Data obtained from 10-day-old pups, including serum hormone concentrations and adrenal expression of steroidogenic enzymes, were analysed using a two-way ANOVA with treatment (vehicle vs FAD286) and sex (male vs female) as between-subject factors. Behavioural data obtained after weaning in the open-field and elevated plus-maze tests, as well as from the salt preference test, were also analysed by two-way ANOVA with treatment and sex as the main factors. When a significant interaction was detected, Tukey’s post hoc test was used for multiple comparisons. Data on serum hormone concentrations and gene-expression data from animals on PND46 were analysed using a three-way ANOVA, with early postnatal treatment, sex, and acute stress condition (non-stressed vs restraint stress) as between-subject factors. Significant interactions were followed by Tukey’s post hoc test, where appropriate. Results are presented as mean ± SD, with individual values shown where applicable. The level of statistical significance was set at p < 0.05. Data analysis was performed using Statistica 9 (StatSoft, Inc., USA).

## 3. Results

### 3.1. Surface righting reflex

The surface righting reflex was present in all pups on PND6, irrespective of treatment or sex. As all animals exhibited the response immediately, latency could not be meaningfully evaluated, and no statistical comparison was performed (data not shown).

### 3.2. Adrenal steroidogenic gene expression and serum hormone concentrations in 10- day-old pups

Early postnatal FAD286 treatment increased adrenal mRNA expression of both *Cyp11b2*, encoding aldosterone synthase, and *Cyp11b1*, encoding steroid 11β-hydroxylase, in 10-day-old pups. Two-way ANOVA revealed a significant main effect of treatment on *Cyp11b2* expression [F(1,19) = 5.23, p = 0.03], with higher expression in FAD286-treated pups than in vehicle-treated controls (Fig. 2A). The main effect of sex [F(1,19) = 0.06, p = 0.81] and the treatment × sex interaction [F(1,19) = 0.71, p = 0.41] were not significant. Similarly, *Cyp11b1* expression was higher in FAD286-treated pups, as indicated by a significant main effect of treatment [F(1,20) = 4.61, p = 0.04] (Fig. 2B). The main effect of sex [F(1,20) = 0.52, p = 0.47] and the treatment × sex interaction [F(1,20) = 0.16, p = 0.69] were not significant.

**Figure 2.**
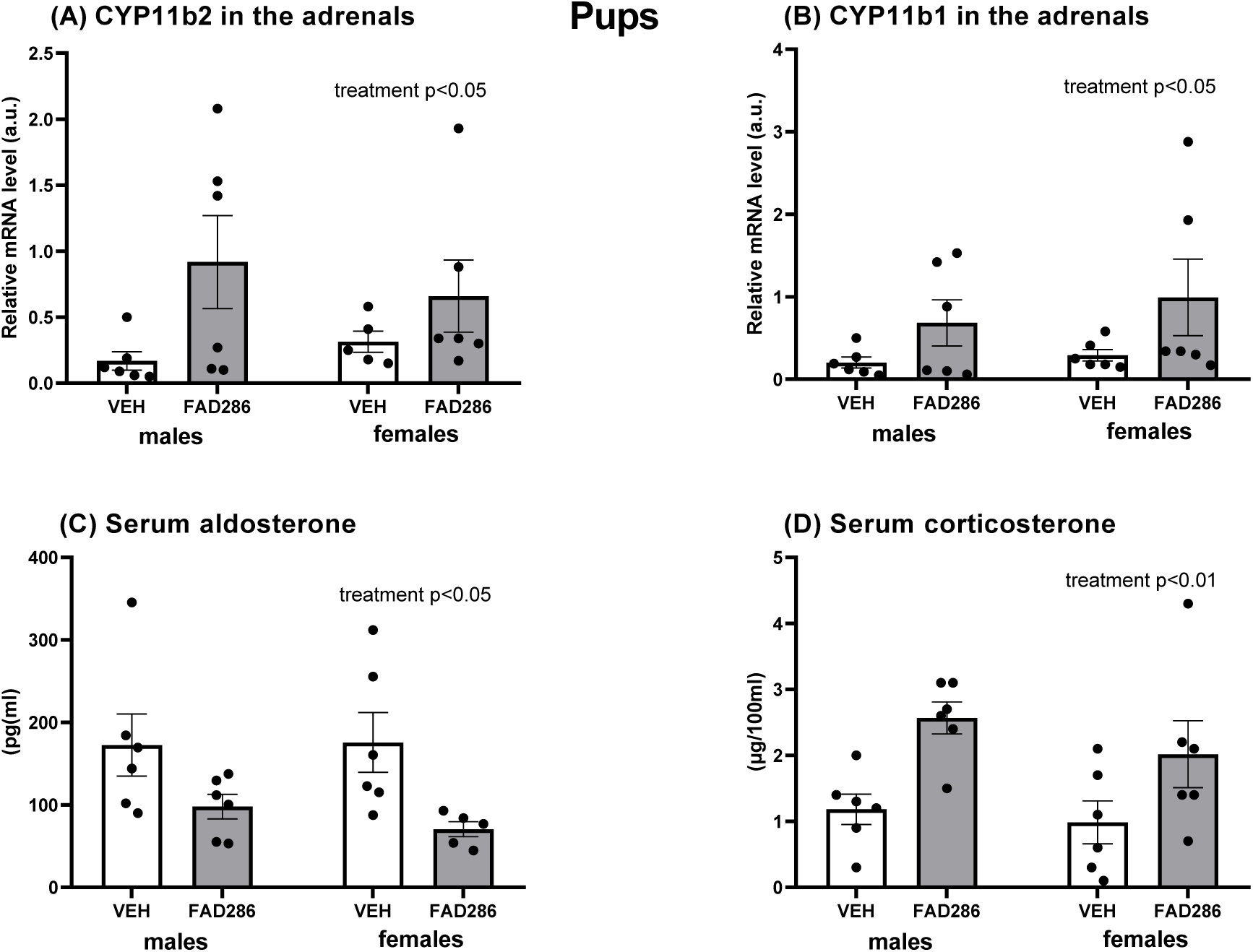
Effects of early postnatal FAD286 treatment on adrenal steroidogenic gene expression and circulating adrenocortical hormone concentrations in 10-day-old male and female rat pups. Adrenal mRNA expression of (A) aldosterone synthase (*Cyp11b2*) and (B) steroid 11β-hydroxylase (*Cyp11b1*), and serum concentrations of (C) aldosterone and (D) corticosterone in male and female pups treated with vehicle (VEH) or FAD286 from PND3 to PND9. Data were analysed using a two-way ANOVA with the main factors sex and treatment. Data are presented as mean ± SD, with individual values shown; n = 5–6 per group. a.u., arbitrary units.

FAD286 treatment also altered circulating adrenocortical hormone concentrations in the pups. Serum aldosterone concentrations were lower in FAD286-treated pups than in vehicle- treated controls [main effect of treatment: F(1,19) = 9.71, p = 0.006] (Fig. 2C). No significant effects of sex [F(1,19) = 0.18, p = 0.67] or treatment × sex interaction [F(1,19) = 0.28, p = 0.60] were observed. In contrast, serum corticosterone concentrations were higher in FAD286-treated pups than in vehicle-treated controls, as demonstrated by a significant main effect of treatment [F(1,20) = 12.29, p = 0.002] (Fig. 2D). The main effect of sex [F(1,20) = 1.18, p = 0.29] and the treatment × sex interaction [F(1,20) = 0.26, p = 0.62] were not significant.

### 3.3. Open field test

Locomotor activity and anxiety-like behaviour were assessed in the open-field test on PND23. Total distance travelled during the open-field test was not significantly affected by early postnatal FAD286 treatment [main effect of treatment: F(1,52) = 1.64, p = 0.20] (Fig. 3A). Two-way ANOVA revealed a significant main effect of sex [F(1,52) = 7.93, p = 0.007], with females travelling a greater distance than males. The treatment × sex interaction was not significant [F(1,52) = 1.45, p = 0.24]. The number of entries into the central area was lower in FAD286- treated rats than in vehicle-treated controls, as indicated by a significant main effect of treatment [F(1,52) = 6.15, p = 0.02] (Fig. 3B). The main effect of sex [F(1,52) = 0.001, p = 0.97] and the treatment × sex interaction [F(1,52) = 0.01, p = 0.91] were not significant. FAD286-treated rats also spent a lower percentage of time in the central area than vehicle-treated controls [main effect of treatment: F(1,52) = 4.01, p = 0.05] (Fig. 3C). No significant main effect of sex [F(1,52) = 0.64, p = 0.42] or treatment × sex interaction [F(1,52) = 0.77, p = 0.38] was observed. Latency to the first entry into the central area was not influenced by treatment [F(1,52) = 2.16, p = 0.14], sex [F(1,52) = 0.63, p = 0.43], or the treatment × sex interaction [F(1,52) = 0.06, p = 0.81] (Fig. 3D).

**Figure 3.**
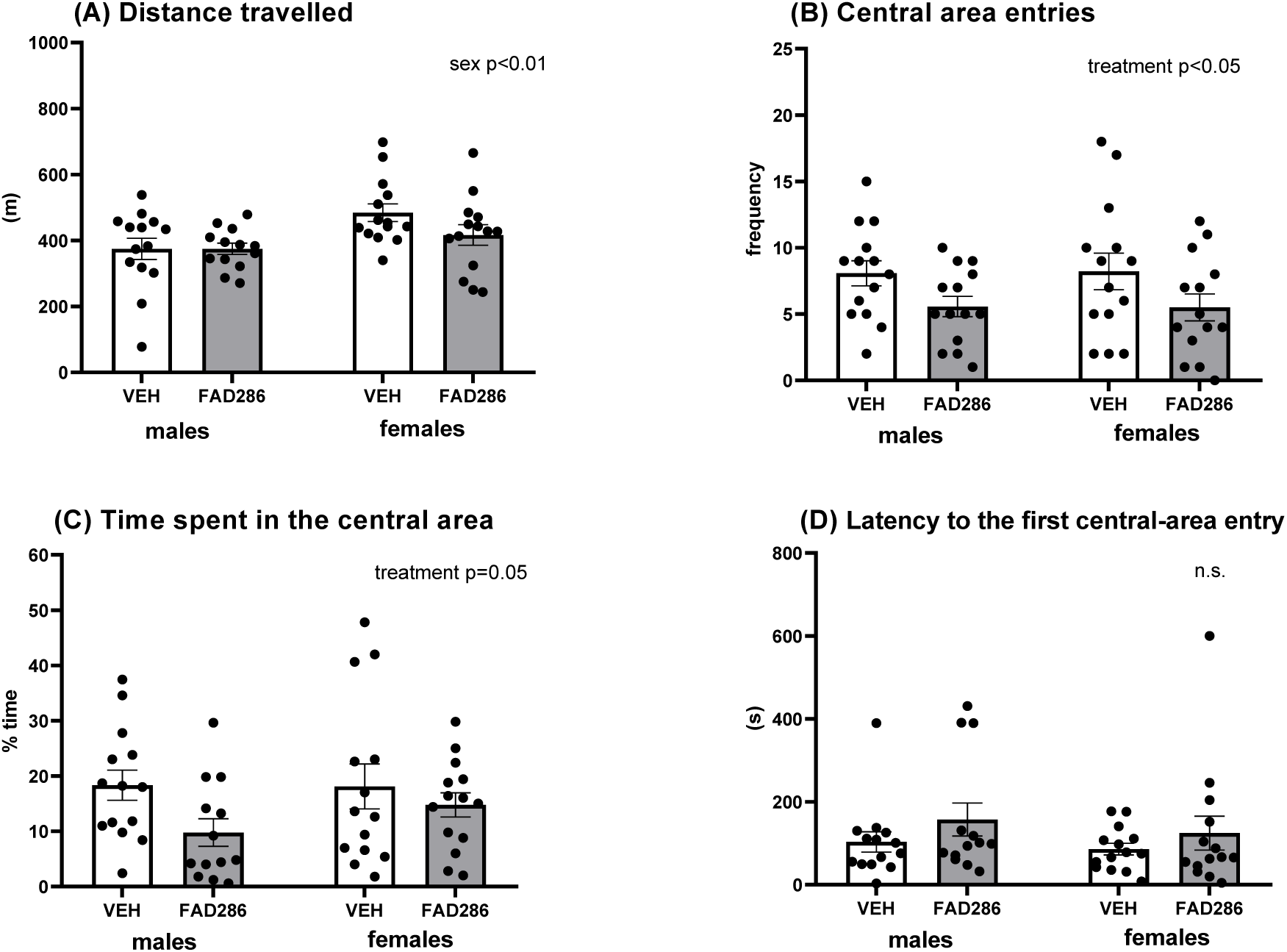
Effects of early postnatal FAD286 treatment on locomotor activity and central-area exploration in the open-field test in male and female rats. Total distance travelled (A), number of entries into the central area (B), percentage of time spent in the central area (C), and latency to the first central-area entry (D) were evaluated on PND23 in male and female rats treated with vehicle (VEH) or FAD286 from PND3 to PND9. Data were analysed using a two-way ANOVA with the main factors sex and treatment. Data are presented as mean ± SD, with individual values shown; n = 14 per group.

### 3.4. Salt preference test

Salt preference and relative NaCl solution intake were analysed using two-way ANOVA, with treatment and sex as between-subject factors. Early postnatal FAD286 treatment did not affect salt preference during the two-bottle choice test. Mean salt preference, calculated from the two consecutive 24-h measurements, was 23.29 ± 6.18% in VEH-treated males and 23.32 ± 6.20% in FAD286-treated males. Corresponding values in females were 22.67 ± 5.58% and 23.15 ± 5.48%, respectively. Two-way ANOVA revealed no significant main effects of treatment or sex and no treatment × sex interaction (all p > 0.05).

Relative intake of the 0.3 M NaCl solution was also unaffected by early postnatal FAD286 treatment. Mean values were 6.81 ± 2.53 and 6.73 ± 2.49 g/100 g body weight/day in VEH- and FAD286-treated males, respectively, and 6.57 ± 2.32 and 6.73 ± 2.49 g/100 g body weight/day in the corresponding female groups. No significant main effects of treatment or sex and no treatment × sex interaction were observed (all p > 0.05).

### 3.5 Elevated plus maze test

Anxiety-like behaviour and ethological measures of risk assessment were evaluated in the elevated plus-maze test on PND29. Two-way ANOVA revealed significant main effects of sex [F(1,52) = 5.76, p = 0.02] and treatment [F(1,52) = 8.04, p = 0.006], as well as a significant treatment × sex interaction [F(1,52) = 9.33, p = 0.003], on the number of open-arm entries (Fig. 4A). Overall, females entered the open arms more frequently than males. Post hoc analysis showed that FAD286 treatment reduced the number of open-arm entries in females (p = 0.001), whereas no significant treatment effect was observed in males.

**Figure 4.**
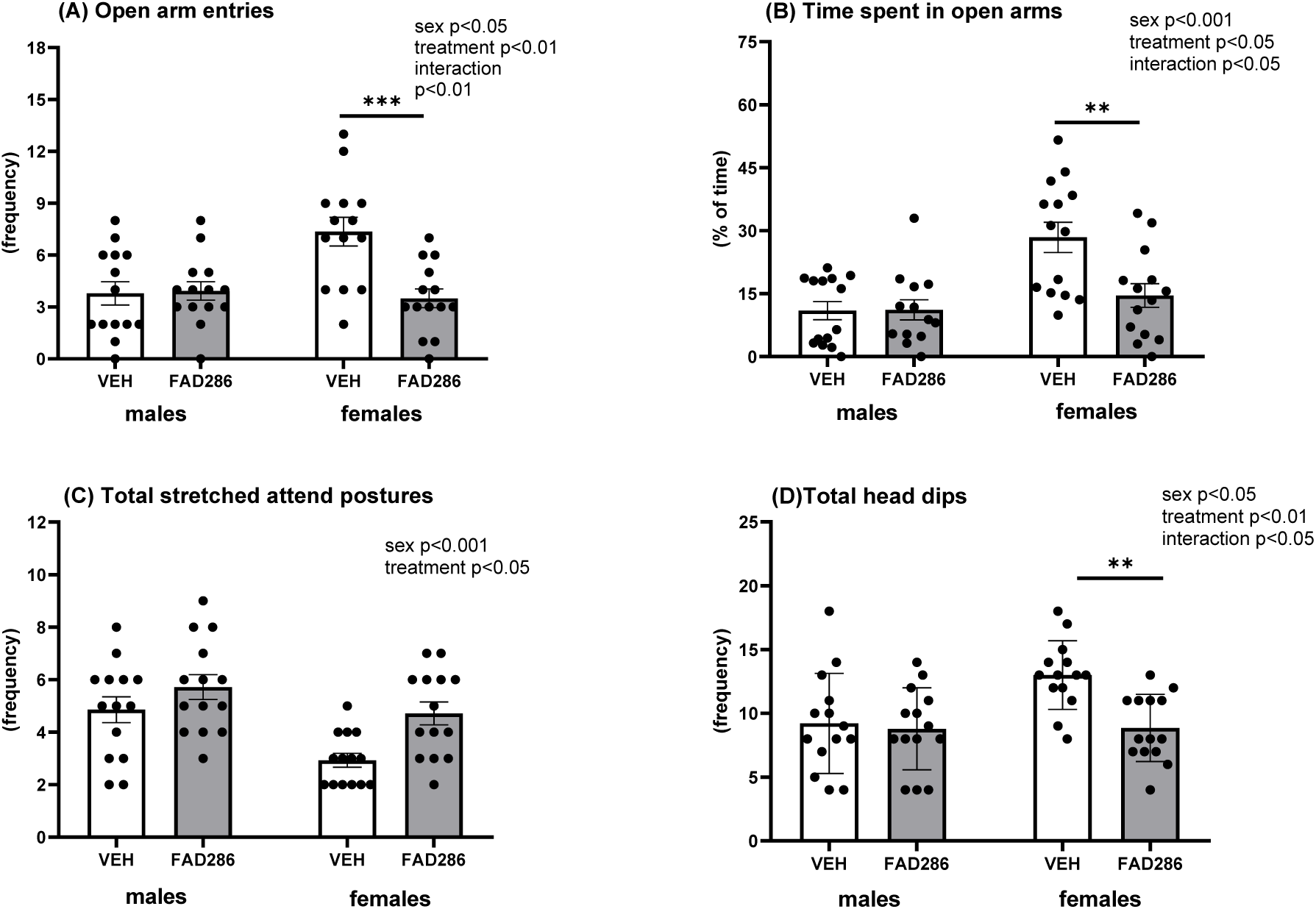
Effects of early postnatal FAD286 treatment on anxiety-like and risk-assessment behaviour in the elevated plus-maze in male and female rats. The number of open-arm entries (A), percentage of time spent in the open arms (B), total number of stretched-attend postures (C), and total number of head dips (D) were evaluated on PND29 in male and female rats treated with vehicle (VEH) or FAD286 from PND3 to PND9. Data were analysed using a two-way ANOVA with the main factors sex and treatment, followed by Tukey post hoc comparisons. Data are presented as mean ± SD, with individual values shown; n = 14 per group. **p < 0.01 and ***p < 0.001 versus the corresponding VEH-treated group.

A similar pattern was found for the percentage of time spent in the open arms (Fig. 4B). Significant main effects of sex [F(1,52) = 13.70, p = 0.0005] and treatment [F(1,52) = 5.78, p = 0.02], together with a significant treatment × sex interaction [F(1,52) = 6.21, p = 0.01], were observed. Females spent a greater percentage of time in the open arms than males. FAD286 treatment significantly reduced the time spent in the open arms in females (p = 0.005), whereas no significant difference between vehicle- and FAD286-treated males was detected.

The total number of stretched-attend postures was also influenced by early postnatal FAD286 treatment (Fig. 4C). Two-way ANOVA revealed significant main effects of sex [F(1,52) = 11.79, p = 0.001] and treatment [F(1,52) = 9.60, p = 0.003], whereas the treatment × sex interaction was not significant [F(1,52) = 1.18, p = 0.28]. Males displayed more stretched-attend postures than females, and FAD286-treated rats exhibited more stretched-attend postures than vehicle-treated animals.

Two-way ANOVA revealed significant main effects of sex [F(1,52) = 5.22, p = 0.03] and treatment [F(1,52) = 7.33, p = 0.01], as well as a significant treatment × sex interaction [F(1,52) = 4.84, p = 0.03] (Fig. 4D) on total head dips. Overall, females displayed more head dips than males. Post hoc analysis showed that FAD286 treatment reduced the number of head dips in females (p = 0.006), whereas no significant treatment effect was observed in males.

The number of closed-arm entries was not significantly affected by treatment [F(1,52) = 0.01, p = 0.91], and the treatment × sex interaction was not significant [F(1,52) = 0.15, p = 0.69]. However, a significant main effect of sex was observed [F(1,52) = 7.71, p = 0.008], with females making more closed-arm entries than males (data not shown).

### 3.6. Adrenal steroidogenic gene expression and adrenocortical hormone responses to restraint stress

To evaluate the persistent effects of early postnatal FAD286 treatment on adrenal steroidogenic regulation and adrenocortical stress responsiveness, adrenal *Cyp11b2* and *Cyp11b1* mRNA expression and serum aldosterone and corticosterone concentrations were determined on PND46 in non-stressed rats and immediately after 120 min of restraint stress.

Adrenal *Cyp11b2* mRNA expression was significantly affected by early postnatal treatment and restraint stress. Three-way ANOVA revealed significant main effects of treatment [F(1,47) = 4.25, p = 0.045] and stress [F(1,47) = 5.49, p = 0.023]. These effects were accompanied by a significant sex × treatment interaction [F(1,47) = 4.40, p = 0.041] and a significant sex × treatment × stress interaction [F(1,47) = 5.27, p = 0.026]. Tukey’s post hoc test showed that, in males, restraint stress increased *Cyp11b2* mRNA expression in FAD286-treated rats compared to their corresponding non-stressed controls (p = 0.012). In restraint-stressed males, *Cyp11b2* expression was also higher in FAD286-treated rats than in VEH-treated rats (p = 0.005). No significant effects of treatment or restraint stress were observed in females. In restraint-stressed FAD286-treated rats, *Cyp11b2* expression was higher in males than in females (p = 0.036). The main effect of sex [F(1,47) = 1.46, p = 0.233], sex × stress interaction [F(1,47) = 1.43, p = 0.238], and treatment × stress interaction [F(1,47) = 3.37, p = 0.073] were not significant (data not shown).

Adrenal *Cyp11b1* mRNA expression was significantly affected by sex [F(1,47) = 42.01, p = 0.001], with higher expression in females than in males. Neither early postnatal treatment nor restraint stress significantly affected *Cyp11b1* expression, and no significant interactions between the factors were observed (all p > 0.05; data not shown).

Three-way ANOVA revealed significant main effects of treatment [F(1,47) = 7.37, p = 0.009] and restraint stress [F(1,47) = 35.43, p < 0.001] on serum aldosterone concentrations (Fig. 5A). Overall, serum aldosterone concentrations were higher in rats treated with FAD286 during early postnatal development than in VEH-treated rats and were increased in restraint-stressed rats compared to the control animals. The treatment × stress interaction was not significant [F(1,47) = 2.99, p = 0.090]. The main effect of sex was not significant [F(1,47) = 0.80, p = 0.375]. Similarly, the sex × FAD286 treatment [F(1,47) = 0.26, p = 0.614], sex × restraint stress [F(1,47) = 0.80, p = 0.377], and sex × FAD286 treatment × restraint stress interactions [F(1,47) = 0.11, p = 0.743] were not significant.

**Figure 5.**
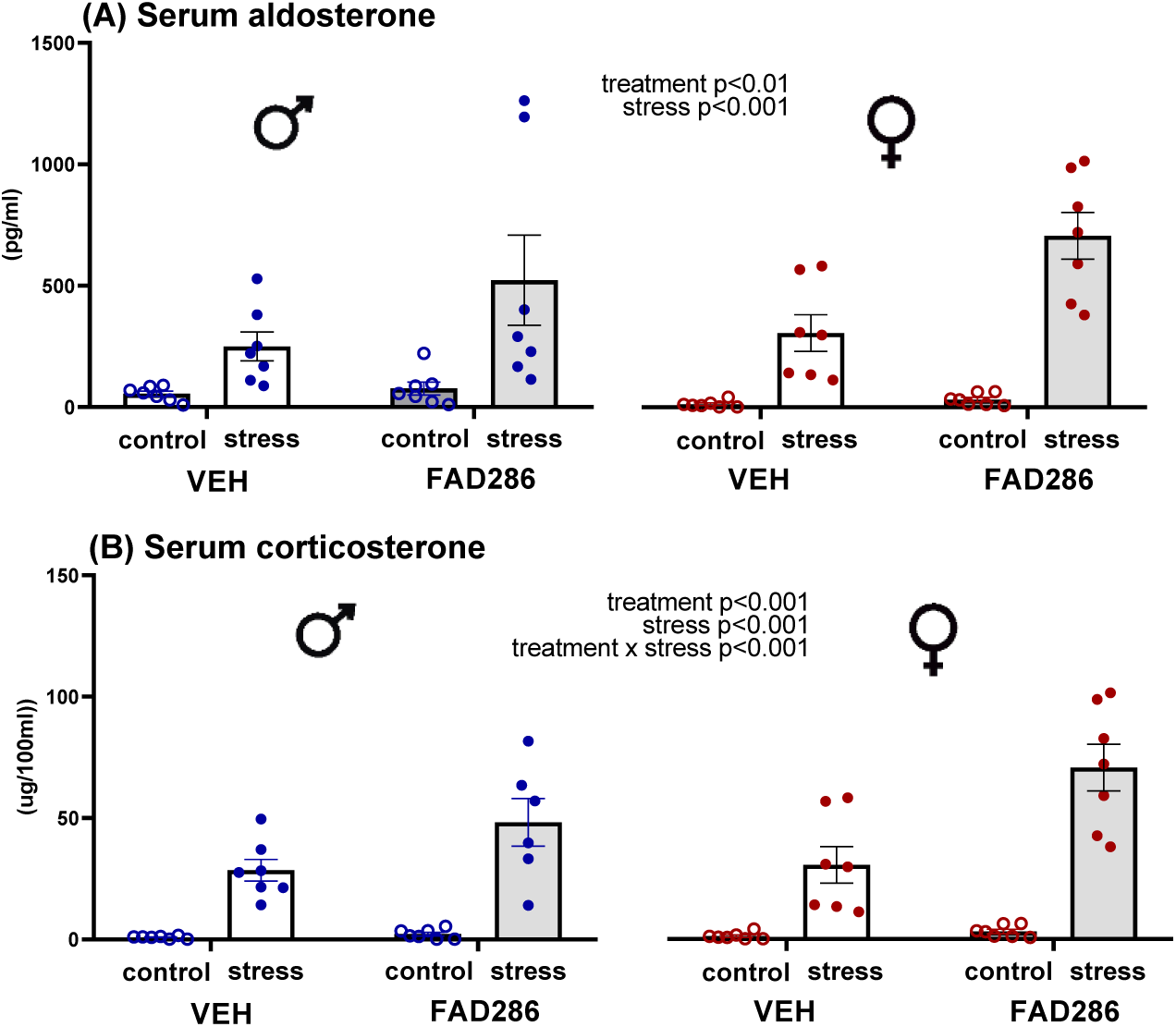
Effects of early postnatal FAD286 treatment on serum aldosterone and corticosterone concentrations in response to restraint stress in male and female rats. Serum concentrations of aldosterone (A) and corticosterone (B) were measured on PND46 in non-stressed control rats and in rats exposed to 120 min of restraint stress. Rats had been treated with vehicle (VEH) or FAD286 from PND3 to PND9. Data obtained from male rats are shown on the left, and data obtained from female rats on the right. Data were analysed using a three-way ANOVA with the main factors sex, treatment and stress. Data are presented as mean ± SD, with individual values shown; n = 6–7 per group.

Serum corticosterone concentrations (Fig. 5B) were significantly influenced by both early postnatal treatment [F(1,47) = 14.7, p < 0.001] and restraint stress [F(1,47) = 111.5, p < 0.001]. In addition, a significant treatment × stress interaction was also observed [F(1,47) = 11.7, p = 0.001], indicating that the effect of early postnatal FAD286 treatment differed between control and stress-exposed rats. Tukey’s post hoc test showed that restraint stress increased serum corticosterone concentrations in both VEH- and FAD286-treated rats (p = 0.001 and p = 0.001, respectively) compared to their corresponding control groups. Following restraint stress, serum corticosterone concentrations increased significantly in rats treated with FAD286 compared to VEH-treated rats (p = 0.001), whereas no difference between the treatment groups was observed in control animals. The main effect of sex was not significant [F(1,47) = 2.9, p = 0.093]. Similarly, the sex × FAD286 treatment [F(1,47) = 2.0, p = 0.165], sex × restraint stress [F(1,47) = 2.4, p = 0.128], and sex × FAD286 treatment × restraint stress interactions [F(1,47) = 1.8, p = 0.187] were not significant.

## 4. Discussion

The present study demonstrates that transient inhibition of aldosterone synthesis during the SHRP led to alterations in anxiety-related behaviour and adrenocortical regulation later in development, with some behavioural effects being sex-dependent. Treatment with the selective inhibitor of aldosterone synthase, FAD286, induced the expected decrease in circulating aldosterone in 10-day-old pups; however, this decrease was accompanied by increased adrenal expression of both steroidogenic genes and elevated serum corticosterone concentrations. During adolescence, FAD286-treated rats showed increased anxiety-like and altered risk-assessment behaviour, with several effects being most pronounced in females. Early FAD286 treatment also resulted in higher overall aldosterone concentrations and an enhanced corticosterone response to restraint stress. Thus, the consequences of transient aldosterone synthase inhibition persisted beyond the treatment period and into later development.

The decrease in aldosterone concentrations on PND10 confirms the expected pharmacodynamic action of FAD286 on aldosterone synthesis. The concomitant increase in Cyp11b2 gene expression may reflect a compensatory response to reduced hormonal output. Because FAD286 inhibits CYP11B2 enzymatic activity rather than *Cyp11b2* transcription, reduced circulating aldosterone can coexist with increased *Cyp11b2* mRNA. This transcriptional response may reflect compensatory upregulation of the inhibited pathway. However, without measurements of renin activity, potassium, ACTH, or CYP11B2 protein and enzymatic activity, its regulatory basis and functional significance remain unclear (Hattangady et al., 2012).

By contrast, the increases in corticosterone concentrations and *Cyp11b1* expression were unexpected findings. In adult rodent models, FAD286 generally leaves corticosterone unchanged at doses that effectively suppress aldosterone, whereas corticosterone decreases at higher exposures, when inhibition of CYP11B1 becomes more relevant (Rigel et al., 2010; Hofmann et al., 2017). Increased corticosterone levels have not been reported as a typical pharmacodynamic effect of FAD286 in these models. The parallel increase in *Cyp11b1* expression in the present study, therefore, argues against direct inhibition of CYP11B1 and is more consistent with activation of the corticosterone-producing pathway. Increased ACTH drive, enhanced adrenal sensitivity to ACTH, or another compensatory change in steroidogenic regulation could contribute to this response.

The developmental stage at which this endocrine response occurred is likely to be important. During the SHRP, low corticosterone exposure is considered part of the protective endocrine environment of the developing brain, whereas aldosterone secretion remains responsive to acute challenges (Sapolsky and Meaney, 1986; Levine, 2001; Varga et al., 2013). FAD286 did not simply lower one adrenal steroid but simultaneously reduced aldosterone and increased corticosterone secretion. This combined response indicates a broader disturbance of the adrenocortical steroid environment during a developmentally sensitive period. Experimental elevation or prolongation of glucocorticoid exposure during gestation or early postnatal life has been associated with later changes in emotional behaviour and stress-axis function, often in a sex-dependent manner (Brummelte et al., 2012; Barbosa Neto et al., 2012). Although these models differ substantially from selective aldosterone synthase inhibition, they demonstrate the sensitivity of later stress regulation to disturbances of corticosteroid exposure during early development. The present data, however, cannot determine which component of the early endocrine change was primarily responsible for the later phenotype.

The behavioural effects of FAD286 were not attributable to impaired early motor development or reduced locomotion, as surface righting, total distance travelled in the open field, and closed-arm entries in the elevated plus-maze were unaffected. Treatment with FAD286 nevertheless reduced central-area exploration and increased stretched-attend postures in both sexes, while reductions in open-arm exploration and head dipping were observed only in females. This pattern indicates altered anxiety-related and risk-assessment behaviour rather than nonspecific behavioural suppression. We previously demonstrated that prolonged aldosterone administration in adult rats similarly increased anxiety-like behaviour without reducing locomotor activity (Hlavacova and Jezova, 2008). Conversely, chronic treatment with the mineralocorticoid receptor antagonist eplerenone produced an anxiolytic profile (Hlavacova et al., 2010). An anxiolytic-like effect has also been observed following hippocampal mineralocorticoid receptor blockade (Bitran et al., 1998).

Although the anxiogenic-like phenotype following early aldosterone synthase inhibition may appear inconsistent with these adult findings, the experimental conditions are not physiological opposites. Aldosterone administration or mineralocorticoid receptor blockade in adulthood acts on a mature corticosteroid system, whereas FAD286 in this study was administered during neuroendocrine development. The latter phenotype may therefore reflect the altered corticosteroid environment during development rather than reduced aldosterone availability alone. The restriction of some behavioural effects to females may reflect sex-related differences in the maturation of corticosteroid-sensitive neural systems. Early-life alterations in glucocorticoid exposure have likewise been associated with sex-dependent behavioural consequences (Brummelte et al., 2012). Periodic maternal deprivation during the SHRP also caused long-term changes that differed between male and female rats (Wigger and Neumann, 1999).

Salt preference was unaffected in FAD286-treated rats, providing no behavioural evidence of a persistent sodium deficit or increased sodium appetite later in development. Voluntary intake, however, is only an indirect indicator of mineralocorticoid homeostasis, and normal salt preference does not exclude changes in renin activity, electrolyte handling or adrenal sensitivity. This possibility remains relevant because sustained FAD286 treatment can activate renin release under dietary conditions that stimulate aldosterone synthesis (Ménard et al., 2006).

The higher aldosterone concentrations on PND46 may indicate a compensatory change in aldosterone regulation following neonatal enzyme inhibition. In adult rats, four weeks of FAD286 treatment increased plasma renin activity under low-sodium conditions, demonstrating activation of upstream compensatory mechanisms during sustained aldosterone synthase inhibition (Ménard et al., 2006). Whether similar mechanisms remained altered after the transient neonatal treatment cannot be determined from the present measurements. Nevertheless, the overall elevation of aldosterone across non-stressed and restraint conditions suggests a change in aldosterone regulation rather than enhanced responsiveness to acute stress. The increase in aldosterone induced by restraint itself is consistent with its established release during acute stress (Stier et al., 2004).

By contrast, early FAD286 treatment enhanced the corticosterone response to restraint without affecting concentrations under non-stressed conditions. This pattern indicates altered regulation of stress-induced rather than basal corticosterone secretion. Evidence that disturbances during the SHRP can modify subsequent corticosterone responses comes from maternal deprivation on PND11, which prolonged corticosterone secretion after behavioural testing later in life (Barbosa Neto et al., 2012). Although maternal deprivation differs substantially from selective inhibition of aldosterone synthase, this finding illustrates the sensitivity of later stress regulation to disturbances in the early adrenocortical environment. The accompanying changes in anxiety-related behaviour further support the possibility that the neonatal endocrine disturbance affected later stress regulation.

The adrenal gene-expression changes on PND46 did not closely parallel the circulating hormone profiles. The induction of *Cyp11b2* in restraint-exposed FAD286-treated males was not accompanied by a corresponding selective increase in aldosterone. Such dissociation is plausible because acute hormone output depends on precursor availability and post-transcriptional regulation, whereas *Cyp11b2* transcription contributes mainly to longer-term steroidogenic capacity (Hattangady et al., 2012). The pattern of *Cyp11b1* expression likewise did not directly correspond to circulating corticosterone levels. Thus, transcript abundance measured after 120 min of restraint should not be interpreted as a direct surrogate of hormone secretion at the same time point.

Several limitations should be considered. The present measurements do not identify the primary signal responsible for either the early or the later endocrine changes. ACTH would help to distinguish altered central HPA-axis stimulation from an adrenal-intrinsic response, while renin activity, electrolytes and steroid intermediates would clarify compensatory regulation and precursor flux. In particular, measurement of 11-deoxycorticosterone and 18- hydroxycorticosterone could establish whether FAD286 altered the distribution of steroid intermediates, as reported in adult rats treated with aldosterone synthase inhibitors (Rigel et al., 2010; Hofmann et al., 2017). Furthermore, mRNA expression provides no information on enzyme protein or catalytic activity, and a single post-restraint time point cannot describe the dynamics of hormone release and recovery.

In conclusion, transient inhibition of aldosterone synthesis during the SHRP reduced aldosterone but increased corticosterone concentrations and adrenal steroidogenic gene expression in rat pups. This early endocrine disturbance was followed by anxiety-related behavioural changes, higher overall aldosterone concentrations and an enhanced corticosterone response to restraint later in development. Behavioural and transcriptional effects showed partial sex dependence, indicating that the later expression of an early adrenocortical disturbance differed between males and females. Although the present design does not isolate the relative contributions of reduced aldosterone and increased corticosterone, the findings suggest that aldosterone availability forms part of the developmental adrenocortical environment during the SHRP and that transient disruption of this environment has consequences extending beyond the treatment period.

## 5. Acknowledgements

This study was supported by the Scholarship for Excellent Researchers R2–R4, funded by the European Union – NextGenerationEU through the Recovery and Resilience Plan for Slovakia, project No. 09I03-03-V04-00442/2024/VA.

## 7. Data availability

The data supporting the findings of this study are available from the corresponding author upon reasonable request.

